# Genomes of *Wiebesia* fig wasps reveal the adaptation and codiversification in fig-fig wasp mutualism

**DOI:** 10.1101/2024.10.22.619299

**Authors:** Bai-Wei Lo, Hsin-Fu Lin, Siu-Wah Kong, Wen-Jer Wu, Selina Cai-Ling Wang, Xuemei Lu, Hurng-Yi Wang

**Affiliations:** Institute of Ecology and Evolutionary Biology, National Taiwan University, Taipei, Taiwan; Research Group of Development and Disease, Max Planck Institute for Molecular Genetics, Berlin, Germany; Graduate Institute of Clinical Medicine, College of Medicine, National Taiwan University, Taipei, Taiwan; Department of Entomology, National Taiwan University, Taipei, Taiwan; Beijing Institute of Genomics, Chinese Academy of Sciences, Beijing, China; Kunming Institute of Zoology, Chinese Academy of Sciences, Kunming, China

**Keywords:** *Ficus pumila* var. *awkeotsang*, *Ficus pumila* var. *pumila*, codiversification, coevolution, olfactory receptor, *Wiebesia*

## Abstract

Figs and fig wasps represent one of the most intimate examples of plant-pollinator coevolution. As figs diversified into geographically isolated populations, both figs and fig wasps underwent selective pressures driven by local adaptation and coevolution. *Ficus pumila* comprises two ecologically distinct varieties: the creeping fig (*F. pumila* var. *pumila*), which is widely distributed across the lowlands of East Asia, and the jelly fig (*F. pumila* var. *awkeotsang*), endemic to Taiwan and found at mid-elevations. To elucidate how codiversification with fig hosts influences the evolutionary trajectories of fig wasps, we analyzed the genomes of *Wiebesia* sp. 2 and sp. 3, the respective pollinators of creeping fig and jelly fig. Our demographic analysis indicates that vicariance during the Last Glacial Period facilitated ecological differentiation between these two fig-fig wasp pairs. Through comparative and population genomic analyses, we identified selection signals linked to habitat adaptation, with evolutionary rates corresponding to the life history traits of their host figs. Variations in host preference behavior, chemosensory gene expression, and adaptive duplications in olfactory receptors highlight potential mechanisms for adaptation to host floral scents. These findings collectively underscore how the obligate mutualism between figs and their pollinating wasps allows the ecological traits and habitat preferences of fig hosts to shape the evolutionary pathways of their pollinators, leaving distinct molecular imprints in the fig wasp genomes. This study demonstrates the capacity of tightly intertwined life cycles between plants and pollinators to drive adaptation and diversification.

## Introduction

The interaction between flowering plants and their insect pollinators has contributed to the immense biodiversity of both groups (Dodd et al. 1999; Grimaldi 1999; Sargent 2004; Kay and Sargent 2009; Cruaud et al. 2012). Despite the prevalence of these interactions, our understanding of the genetic processes involved in codivergence at the microevolutionary scale—the initial step in generating biodiversity—remains limited (Hembry et al. 2014; Medina et al. 2022). One challenge in studying codiversification is the complexity of organisms’ interactions with their environments, making it difficult to disentangle the selective pressures from multiple biotic and abiotic sources (Althoff et al. 2014).

The mutualism between pollinating fig wasps (Hymenoptera: Agaondiae) and figs (Moraceae: *Ficus*) offers an exceptional system for exploring coevolution (Janzen 1979; Weiblen 2004; Herre et al. 2008). Figs have evolved peculiar, fruit-like inflorescences called syconia (singular: syconium) that house their flowers. These flowers are exclusively pollinated by fig wasps, which, in turn, rely on specialized gall flowers (short-styled and sterile) within the syconia to lay their eggs. Fig wasps spend almost their entire life cycle within the syconia. The pollen-bearing female wasps only leave the syconia after mating and have just a few days to search for another syconium. Thus, figs and fig wasps share a common abiotic environment, while the biotic environment encountered by the wasps is largely restricted to the community within the fig syconia.

The female fig wasps are attracted to the unique floral volatile compounds of their associated hosts (Ware et al. 1993). Diversification of host floral scents affects their associated fig wasps’ preference behavior, and can contribute to pre-zygotic barrier (Ware et al. 1993; Grison-Pigéet al. 2002; Cornille et al. 2012; Souto-Vilarós et al. 2018). Recognition of scent is thus key to the specificity of the mutualism. The tightly bounded life cycles between figs and pollinating wasps suggest that ecological traits, such as life history, habitat preference, and floral scents of fig host should have profound impact on the fundamental evolutionary process of their associated pollinator, and left molecular signatures in the genomes of pollinators (Yoder 2016).

Recent advances in genomic studies on non-model organisms have illuminated the coevolutionary dynamics between figs and fig wasps (Zhang et al. 2020; Wang et al. 2021). However, few studies have examined how differentiating populations are influenced by divergent coevolutionary pressures (Satler et al. 2019). Consequently, genomic investigations focused on closely related lineages are crucial for understanding the process of fig-fig wasp codiversification.

The varieties of *Ficus pumila* L. and their associated pollinator wasps present ideal candidates for such studies. *Ficus pumila* is a dioecious species that consists of two ecologically differentiated varieties. *Ficus pumila* L. var. *pumila*, known as the creeping fig, is widely distributed in the lowlands of East Asia, while *F. pumila* var. *awkeotsang* (Makino) Corner, or jelly fig, is found in the mid-elevations of Taiwan (Fig. 1a, 1b). These two varieties differ in life history traits, including syconia size and phenology. (Fig. 1a) (Hsieh et al. 1993). Male creeping figs undergo three to four flowering cycles annually, whereas male jelly figs typically have one or two (Shih 1988; Ho et al. 1998). The varieties also differ in their floral volatile compound compositions (Chen and Wu 2010; Chen et al. 2016).

**Fig. 1.**
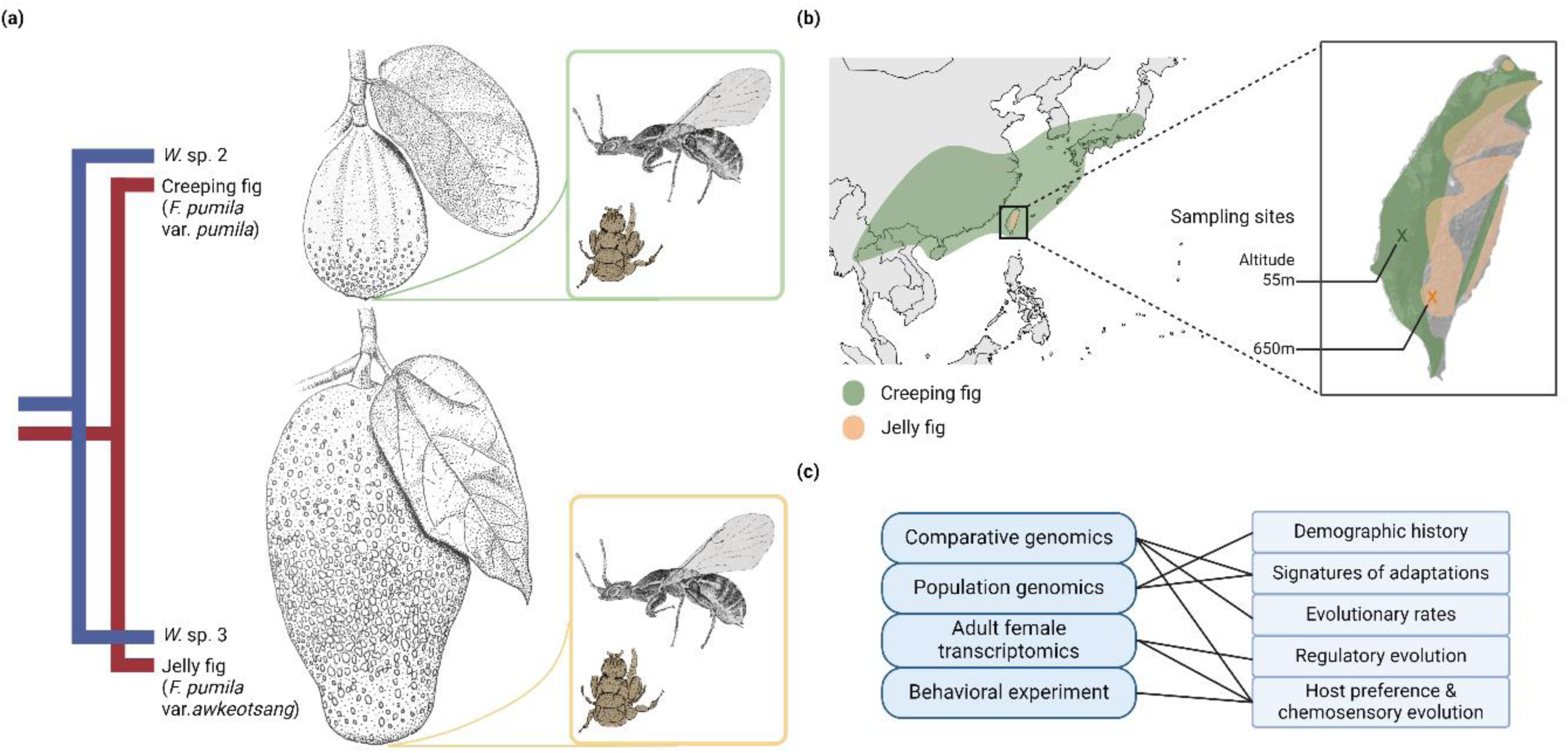
Research system. a) The cophylogeny of the studied *Wiebesia* spp. and their *Ficus pumila* hosts (Wang et al. 2013). Branchlets with syconia of the two fig varieties and their respective fig wasps were illustrated by Bai-Wei Lo. b) The presumed distribution of *Ficus pumila* varieties and the locations where the fig wasp genomic sequencing samples were collected. Geographical distribution of the two *Ficus pumila* varieties was modified from GBIF.org (2024). c) A summary of the experiments and analyses conducted. Created with Biorender.com.

The ecology of *Ficus pumila* pollinators has been less thoroughly studied. Originally classified as a single species, *Wiebesia pumilae*, they have since been split into three species based on genetic evidence (Chen et al. 2012). *Wiebesia* sp. 1 and its southern sibling, *W.* sp. 2, pollinate the creeping fig, whereas *Wiebesia* sp. 3 pollinates the jelly fig. A pioneering genomic study on creeping fig and *W.* sp. 2 from Mainland East Asia has provided important insights into chemosensory coevolution in this pollination system (Wang et al. 2021). However, the biogeographic history and genomic mechanisms underpinning the codiversification between these fig wasps and fig varieties are not yet understood.

In this study, we present the genomic sequences of *W.* sp. 2 and *W.* sp. 3 from Taiwan, the pollinators of the creeping fig and jelly fig, respectively. These genomes offer comprehensive insights into the genomic processes of codiversification. We first reconstruct the demographic history to explain the niche partitioning of the two species. We then explore the genomic signatures of adaptations to different niches. Through behavioral experiments and transcriptomics, we examine the evolution of host preference in these closely related wasps. Finally, by integrating olfactory receptor annotations from other fig wasps, we investigate the consequences and potential genetic basis of host specificity in fig-fig wasp coevolution.

## Results

### Genome sequencing, assembly, and annotation

We *de novo* assembled the genome of *Wiebesia* sp. 3 with PacBio long reads (Table S1), and applied Blobtools to remove possible contaminations. Subsequent polishing of the draft genome using both long and short reads yielded a final genome of 323.8 Mbp (Table 1). It comprises 230 contigs (N50=18.6 Mbp), with a GC content 30.16% and no sign of residual contamination (Fig. S1). The complete BUSCO score of *W.* sp. 3 genome is 93.7%. Ninety-seven percent of the genome sequences were assembled according to kmer estimates. The assembly statistics and BUSCO scores together suggest that this genome is highly contiguous and comparable to other fig wasp genomes (Xiao et al. 2013; Zhang et al. 2020; Wang et al. 2021) (Table S2, Fig. S2). Additionally, we obtained 80x short reads from a *W.* sp. 2 population in Taiwan and performed a reference-based assembly.

**Table 1.**
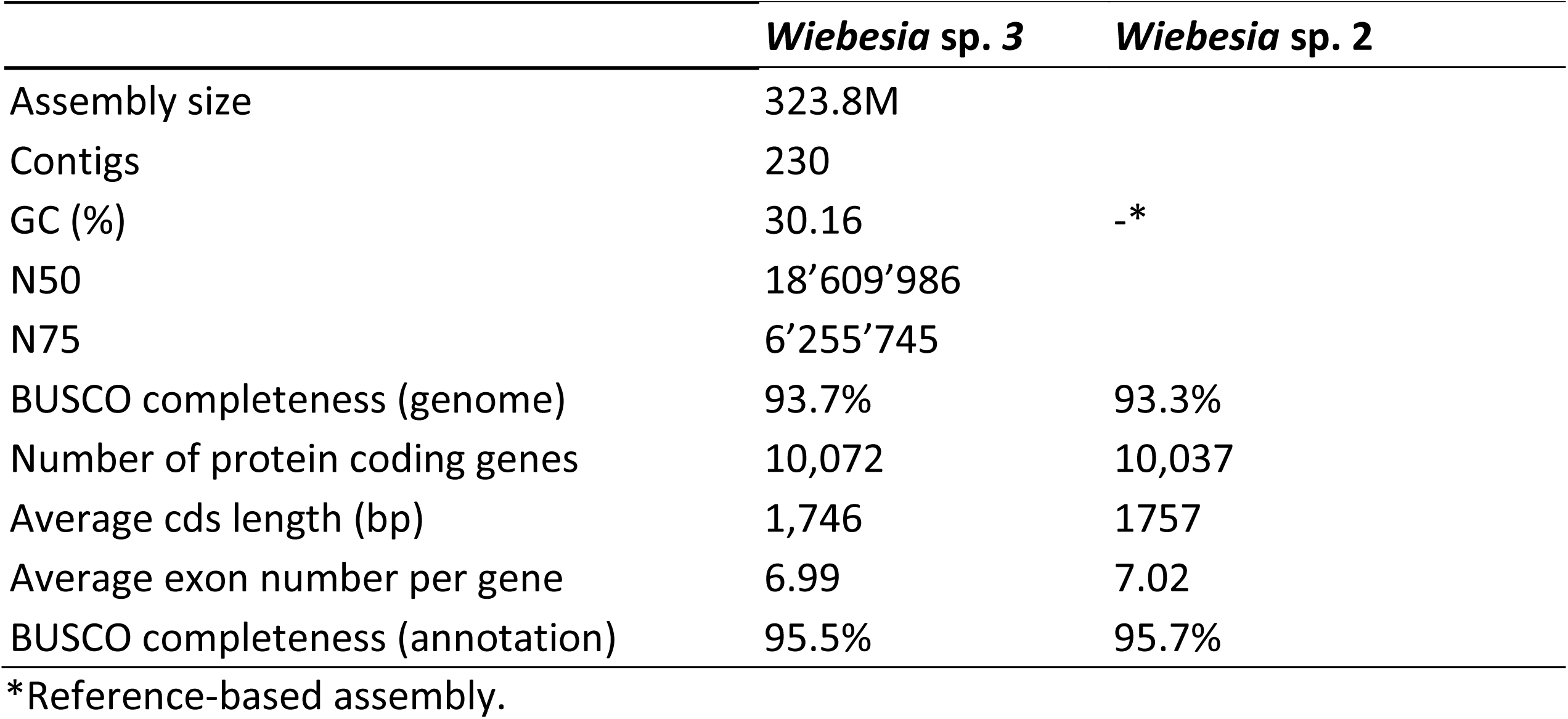
Assembly statistics of two *Wiebesia* species.

We independently annotated the two genomes employing the same protocol. Approximately 10% of repetitive sequences were identified in the genome (Table S3). Using the EVidenceModeler pipeline, 13,342 and 13,684 protein coding genes of *W.* sp. 3 and *W.* sp. 2, were annotated respectively (Table S4). After manual curation with Web Apollo, 10,072 and 10,037 genes were left in the gene sets (Table 1). Despite reduction in gene numbers, BUSCO values of the two species’ gene sets after manual curation improved from 93.0% to 95.5% in *W.* sp. 3 and from 92.8% to 95.7% in *W.* sp. 2. As a result, these annotations are among the most complete of all available fig wasp genomes (Table S2). Blast2GO annotation of functional genes reported that 95% of the genes have at least one match hit against public datasets (Figure S3).

### Species divergence corresponds to the Last Glacial Period

The genomic DNA libraries were constructed from 30 pooled individuals from the respective species. To understand the divergence history of the two species, we applied the PoolParty pipeline to estimate genome-wide population genomic parameters. We then used fastsimcoal2 to simulate and evaluate different demographic models (Table S5). Among all models tested, the most supported scenario is the one considering both early and recent gene flow between *W.* sp. 3 and *W.* sp. 2 (Fig. 2; Table S5). In this scenario, *W.* sp. 3 and *W.* sp. 2 diverged during the Middle Pleistocene, about 550,000 years ago, followed by subsequent gene flow, predominantly from *W.* sp. 3 to *W.* sp. 2. Gene flow ceased from ca. 80,000 until 10,000 years ago. Notably, this complete isolation period overlaps with the Last Glacial Period (115,000–11,700 years ago; Dansgaard et al. 1993). Afterwards, gene flow was reestablished, with more prominent migration from *W.* sp. 2 towards *W.* sp. 3.

**Fig. 2.**
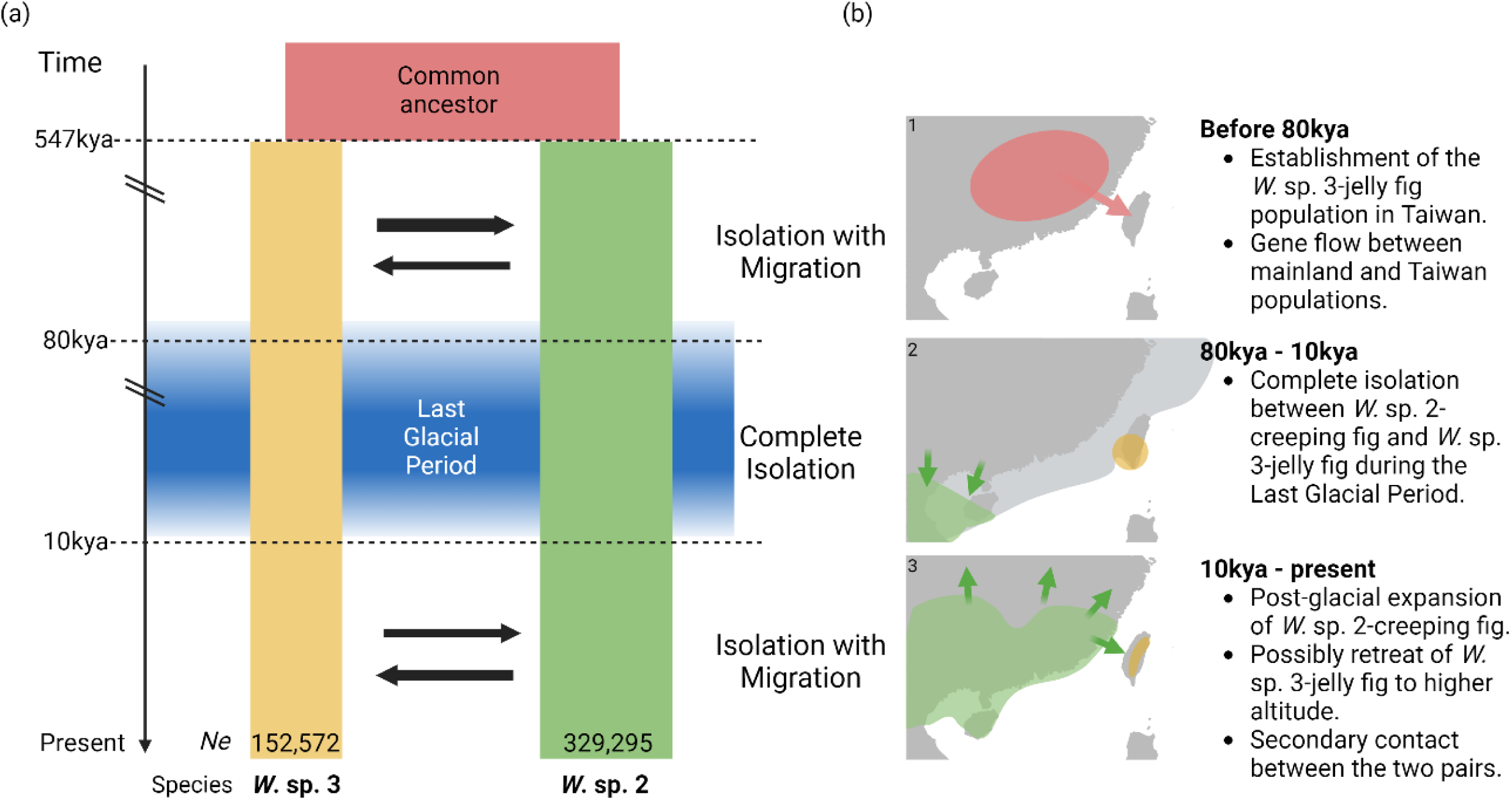
Demographic history of the two *Wiebesia* fig wasp species. a) The demographic model inferred by fastsimcoal2 shows a time of complete isolation that largely overlaps with the Last Glacial Period. The arrows in the middle represent gene flow between species. b) Proposed scenarios corresponding to each time segment based on current genetic and ecological evidence. The distribution of *W.* sp. 2 between 80kya – 10 kya in b2 was based on the warm temperate evergreen forest biome predicted in the Last Glacial Maximum (Harrison et al. 2001). In present day, this biome largely overlaps with the distribution of *W.* sp. 2 (Chen et al. 2012), which is illustrated in b3. Created with BioRender.com.

### Evolutionary rate differences and metabolic-associated adaptations

We compared the evolutionary rates between *W.* sp. 2 and *W.* sp. 3. The non-synonymous substitutions (dN) and the synonymous substitutions (dS) of the *W.* sp. 3 lineage were 0.88 x 10^-3^ and 5.89 x 10^-3^, respectively. These estimates are significantly lower (*p* < 10^-10^) than those (dN: 1.15 x 10^-3^ and dS: 7.05 x 10^-3^) of *W.* sp. 2 (Table 2). We also compared the evolutionary rates of different *W.* sp. 2 populations across the Taiwan Strait using a published *W.* sp. 2 genome from mainland China (Wang et al. 2021). Intriguingly, within *W.* sp. 2, China population has a 6% (dN) and a 4% (dS) higher substitution rate than *W.* sp. 2 from Taiwan (Table 2).

**Table 2.**
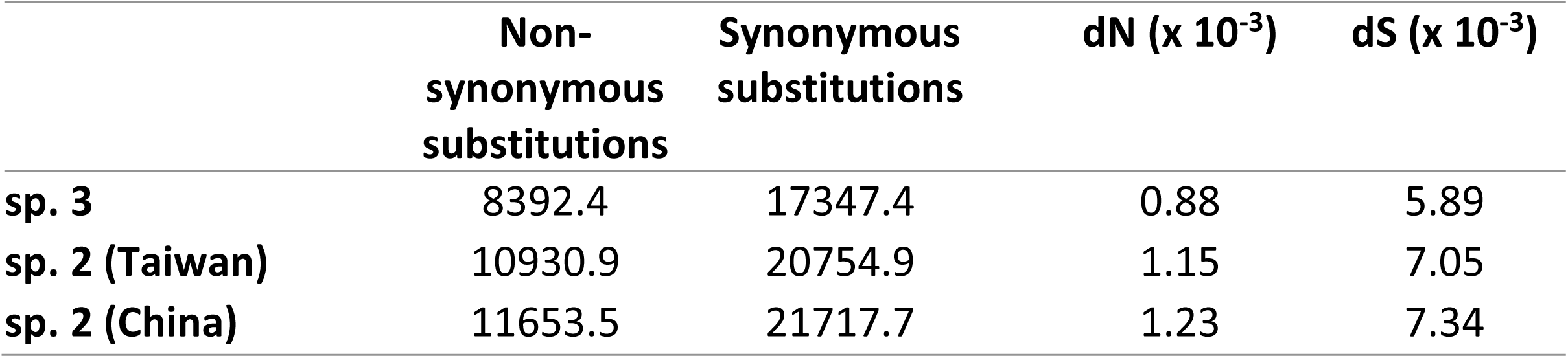
Estimation of nonsynonymous, synonymous substitutions, dN, and dS in each *Wiebesia* population accumulated since the divergence of *W*. sp 2 and *W*. sp. 3.

|  | **Non-synonymous substitutions** | **Synonymous substitutions** | **dN ($\times 10^{-3}$)** | **dS ($\times 10^{-3}$)** |
| --- | --- | --- | --- | --- |
| **sp. 3** | 8392.4 | 17347.4 | 0.88 | 5.89 |
| **sp. 2 (Taiwan)** | 10930.9 | 20754.9 | 1.15 | 7.05 |
| **sp. 2 (China)** | 11653.5 | 21717.7 | 1.23 | 7.34 |

Out of the 9,352 single copy orthologs between the *Wiebesia* species, 643 have signatures of selection (dN/dS ratio > 1), which turned out to be enriched in GOs associated with serine-type endopeptidase and respiratory chain complex IV (Table 3). Tajima’s D test showed that a total of 433 protein-coding genes in the *W.* sp. 3 genome exhibited negative values, implicating population growth or positive selection. GO analysis revealed that these genes are enriched in metabolic process, proteasome complex, and molecular binding (Table S6). In *W.* sp. 2, 348 genes were shown to have negative Tajima’s D values but no specific enriched GO term being identified. In summary, using both direct and linked selection-based methods, we demonstrated that genes putatively under positive selection are enriched in energy production and metabolism.

**Table 3.** Gene ontologies enriched in dN/dS > 1 dataset. All GOs significant at 0.05 FDR level were shown.

| GO term | GO ID | FDR |
| --- | --- | --- |
| **Component** |  |  |
| Respiratory chain complex IV | 0045277 | 0.0031 |
| **Process** |  |  |
| Mitochondrial electron transport, cytochrome c to oxygen | 0006123 | 0.0075 |
| Proteolysis | 0006508 | 7.60E-07 |
| **Function** |  |  |
| Cytochrome-c oxidase activity | 0004129 | 0.0031 |
| Serine-type endopeptidase activity | 0004252 | 9.47E-21 |

### The two *Wiebesia* species are unequally divergent in host specificity

To understand host attraction phenotype and how it affects host specificity in this coevolutionary system, we conducted behavioral experiments on the two *Wiebesia* species. We separately introduced the two species to their alternative host variety’s male and female receptive fig syconia and used their natural host variety as a control. *W.* sp. 3 sporadically entered the male syconia of creeping figs (Table 4). However, it has a notably lower and less consistent entering rate (0.2±0.2) compared to the natural pollinator *W.* sp. 2, which enters at a steadier and higher rate (0.3±0.06, *p* value < 10^-16^). In contrast, *W.* sp. 2 had never entered the male syconia of jelly figs in all 24 repeated experiments. Our behavioral experiments concluded that the two closely related fig wasps are partially host specific and unequally divergent in their host preference behavior.

**Table 4.**
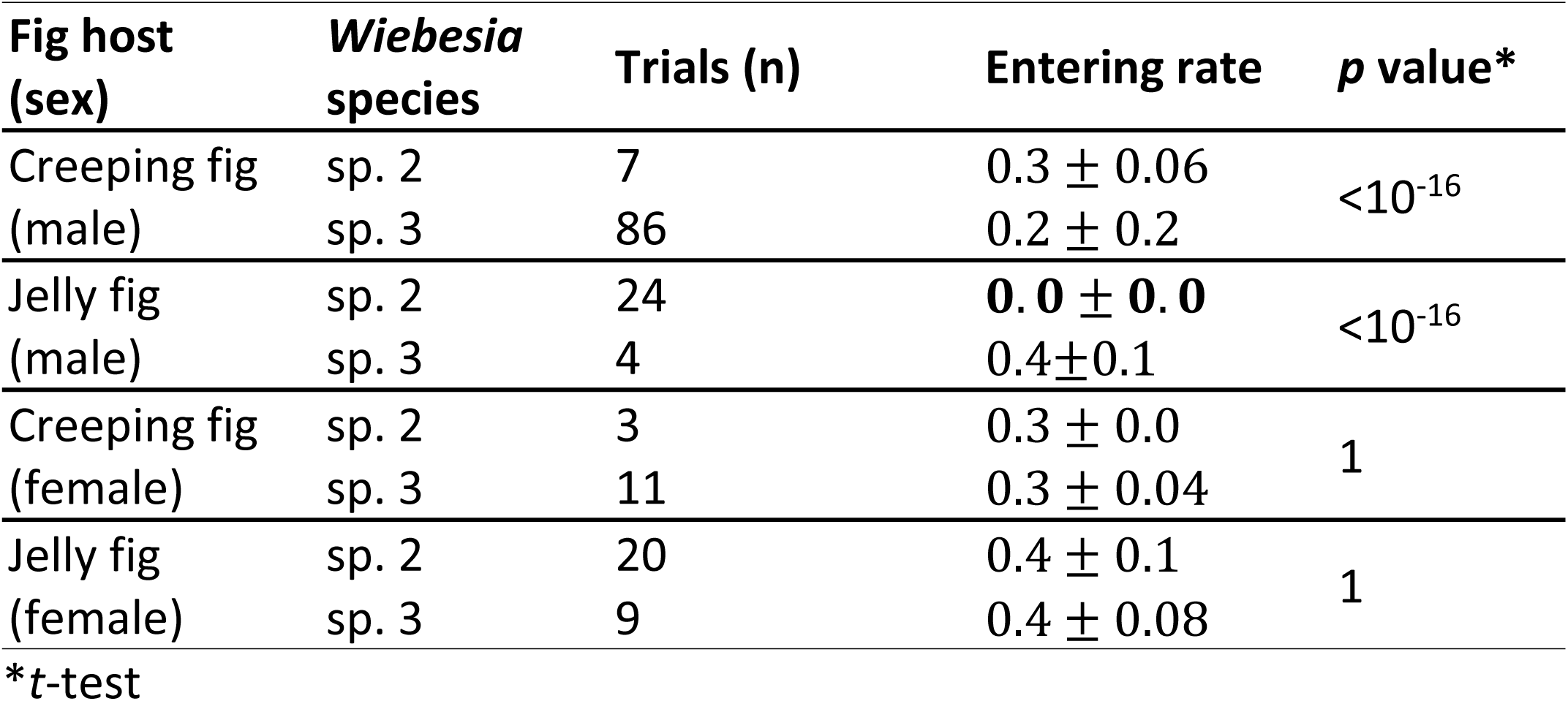
Results of host attraction experiment.

| Fig host (sex) | *Wiebesia* species | Trials (n) | Entering rate | *p* value* |
| --- | --- | --- | --- | --- |
| Creeping fig (male) | sp. 2 | 7 | $0.3 \pm 0.06$ | $<10^{-16}$ |
| | sp. 3 | 86 | $0.2 \pm 0.2$ | |
| Jelly fig (male) | sp. 2 | 24 | **$0.0 \pm 0.0$** | $<10^{-16}$ |
| | sp. 3 | 4 | $0.4 \pm 0.1$ | |
| Creeping fig (female) | sp. 2 | 3 | $0.3 \pm 0.0$ | 1 |
| | sp. 3 | 11 | $0.3 \pm 0.04$ | |
| Jelly fig (female) | sp. 2 | 20 | $0.4 \pm 0.1$ | 1 |
| | sp. 3 | 9 | $0.4 \pm 0.08$ | |

\**t*-test

### Divergence of olfactory expression profiles

To examine if changes in gene regulation lead to divergence in host preferences between the two wasp species, we sequenced and compared the adult female transcriptomic data of *W.* sp. 2 and *W.* sp. 3 (Fig. S4, S5). Among the 8,701 expressed genes, 432 showed differential expression between two species. GO enrichment analysis revealed that the differentially expressed genes are involved in olfaction, cellular signal transduction, serine type endopeptidase activity, and cuticle structure (Table 5). Chemosensory genes, which were differentially expressed in the two *Wiebesia* species, appeared to be single copy in *W.* sp. 2 and *W.* sp. 3, as well as the model fig wasp species *Ceratosolen solmsi*. Notably, *OBP11*, that detects repellent scent signals in *W.* sp. 2 (Wang et al. 2021), was significantly down regulated in *W.* sp. 3 (Table S7). Additionally, among chemosensory genes, olfactory receptors not only showed differential expression between species but also had elevated overall dN/dS ratios, with a significantly higher proportion of genes exhibiting a dN/dS ratio > 1 compared to the genome-wide background (Table S8).

**Table 5.** Gene ontologies enriched in cross-species differential expressed gene set.

| GO term | GO ID | FDR |
| --- | --- | --- |
| **Component** |  |  |
| Extracellular region | 5576 | 8.75E-06 |
| **Process** |  |  |
| Sensory perception of smell | 7608 | 1.66E-05 |
| G protein-coupled receptor signaling pathway, coupled to cyclic nucleotide second messenger | 7187 | 0.0015 |
| **Function** |  |  |
| Serine-type endopeptidase activity | 4252 | 1.66E-05 |
| Ion channel activity | 5216 | 4.74E-05 |
| Olfactory receptor activity | 4984 | 3.72E-04 |
| Odorant binding | 5549 | 4.95E-04 |
| Ion gated channel activity | 22839 | 0.0015 |
| Structural constituent of cuticle | 42302 | 0.0231 |

### Lineage-specific expansions and reductions of olfactory receptor family

To better understand how codivergence with host scents shapes the evolution of olfactory receptor (OR) genes of pollinating fig wasps on a broader evolutionary scale, we expanded our samples to include fig wasp from another dioecious lineage: *C. solmsi*, and two monoecious fig wasp species: *Elisabethiella stuenckenbergi* and *Eupristina verticillata*. Thus, our comparisons covered lineages from different host sexual systems, facilitating insights into OR family size evolution (Fig. 3). While the number of OR repertoires was largely conserved among all studied fig wasps, independent gains and losses of OR repertoires were found in each major lineage of pollinating fig wasps (Fig. 3).

**Fig. 3.**
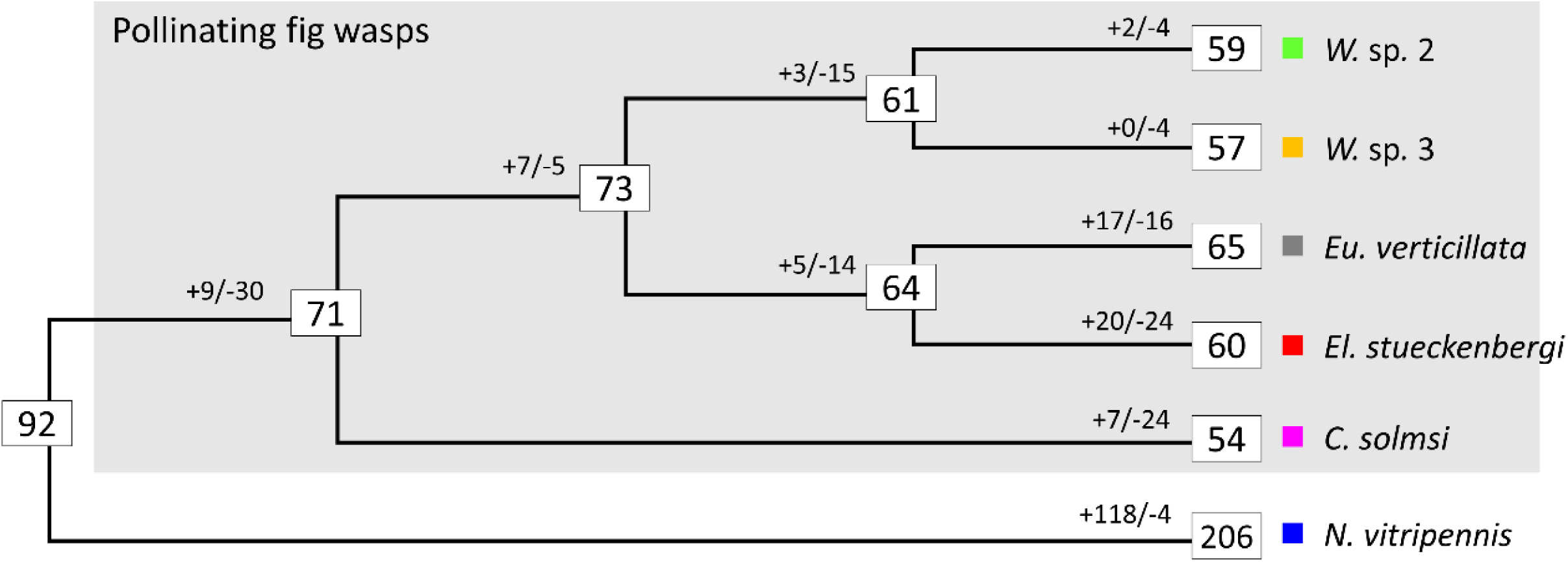
Maximum likelihood phylogeny of studied fig wasps and olfactory receptor (OR) family size evolution. Numbers inside the boxes denote size of ORs at the nodes and leaves. Numbers with a plus or minus sign before the boxes denote gene gain and loss along the branch. Abbreviations: *Eu*, *Eupristina*; *El*, *Elisabethiella*; *C*, *Ceratosolen*; *N*, *Nasonia*.

An in-depth examination of OR gene tree (Fig. S6, Fig 4) revealed details of lineage-specific gene gains and losses. Notably, the subfamily Z has independently expanded in all four studied fig wasp genera (Fig. 4a). Other scattered lineage-specific expansions can be found across the gene tree (eg. Fig. 4c, also see Fig. S6). Further syntenic comparisons indicated that these expansions are due to multiple local tandem repetitive arrays (Fig. 4b & d). Collectively, these data indicate that OR genes in pollinating fig wasps have evolved dynamically, with tandem duplications contributing to the emergence of novel OR genes.

**Fig 4.**
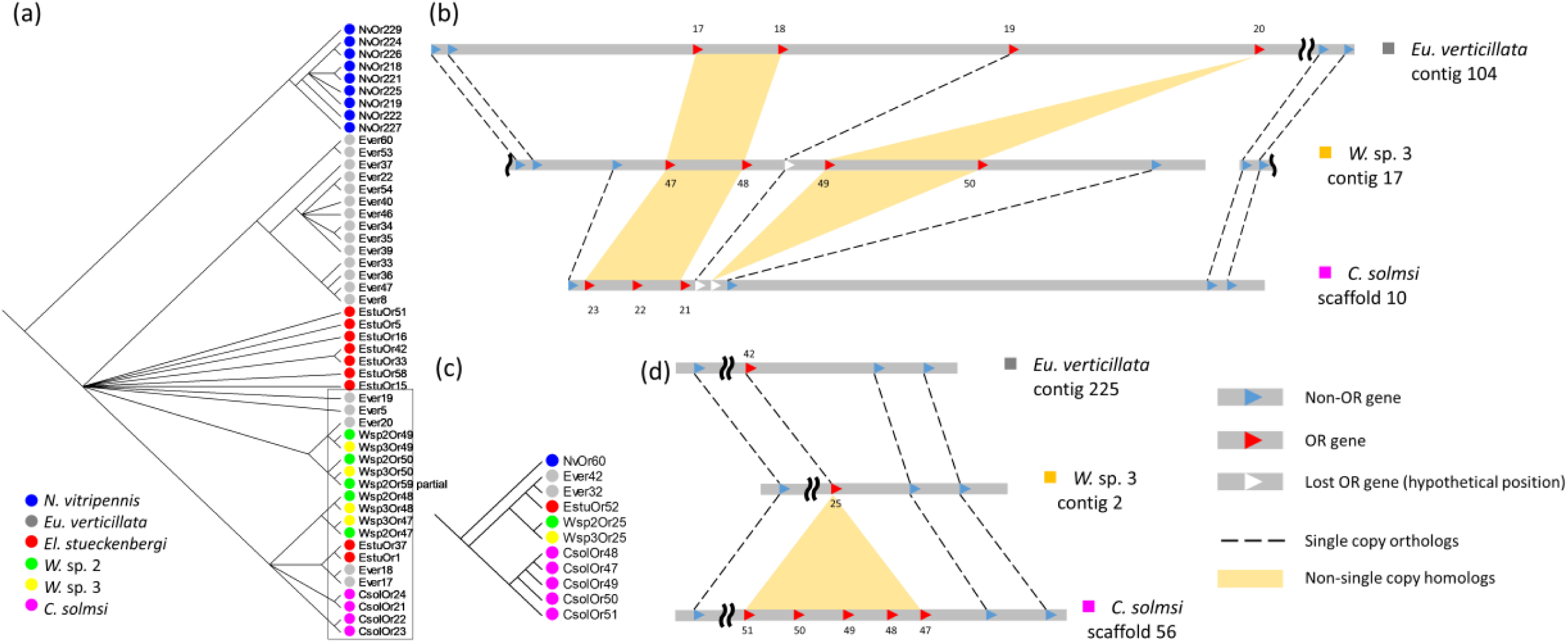
Examples of OR gene expansions due to tandem duplications in fig wasps. a) Subfamily Z of OR. Noting that all studied species experienced gene expansions, including the outgroup parasitoid wasp *Nasonia vitripennis*. The synteny of the clade in square is shown in (b). Nodes with less than 50% bootstrap support were collapsed into polytomy. b) Local synteny of part of subfamily Z shows the dynamic nature of lineage specific gene duplications and losses. *Elisabethiella stueckenbergi* is excluded in the analysis because the OR annotation was derived from transcriptome. c) & d) Tandem duplications resulted in a *Ceratosolen solmsi*-specific expansion.

### Accelerated evolution in duplicated olfactory receptors

To test whether the duplicated ORs experienced different selective pressure, we applied codon-based branch site selection analysis. We first assessed the quality of the annotated ORs by predicting the number of transmembrane domains present in these genes. Only species with an average number of TDs above five were used to ensure adequate information (Table S9). These included *C. solmsi* and the two *Wiebesia* species. The ORs from these three species were divided into two groups: simple orthogroups and complex orthogroups (Fig. S7). The former consist of single copy orthologs from the three species, while the latter contain monophyletic gene trees that include duplicated paralogs. Notably, five out of the 41 branches in complex orthogroups exhibited signatures of positive selection, a rate significantly higher than the two out of 87 branches in simple orthogroups (*p* < 0.05). This indicates a greater prevalence of positive selection in complex orthogroups compared to simple ones, suggesting that duplicated ORs are more likely to undergo positive selection.

## Discussion

In this study, we investigated the codiversification genomic landscape in two closely related *Wiebesia* fig wasp species. By combining genomic, transcriptomic, and behavioral data, we revealed the genetic signatures underlying ecological specialization and olfactory coevolution in these pollinating wasps. Our findings highlight the strong influence of host life histories and ecological conditions on the evolutionary trajectories of their pollinators, demonstrating how tightly linked life cycles shape adaptation and diversification.

### The divergence and ecological differentiation of *Wiebesia* spp. and their fig hosts

Our demographic modeling indicates that the common ancestor of *W.* sp. 2 and *W.* sp. 3 began diverging 547,000 years ago during Middle Pleistocene, likely representing the colonization of the ancestral fig-fig wasp from Mainland East Asia to Taiwan (Fig. 2b1). The estimated population sizes of the two species also reflect differences in their distribution ranges, with the *Ne* of the widely distributed *W.* sp. 2 being about twice that of the endemic *W.* sp. 3 (Fig. 2a). Migration between the two species ceased 80,000 years ago, leading to complete isolation, which coincides with the beginning of the Last Glacial Period (115,000 – 11,700 years ago) (Fig. 2b2). During this time, jelly fig-*W.* sp 3 likely remained confined to Taiwan, while their mainland siblings retreated to warmer southern refuge. This is consistent with the retreat of creeping fig favored-biome in the Last Glacial Maximum based on ancient pollen (Yu et al. 2000; Harrison et al. 2001). Migration resumed around 10,000 years ago and persists to the present day (Fig. 2a). This coincides with the post-glacial expansion of creeping fig in East Asia, which included the recolonization of creeping fig/*W.* sp. 2 to the lowlands of Taiwan (Fig. 2b3) (Wang et al. 2013). Previous studies based on genetic markers have linked Pleistocene climate oscillations to *Wiebesia* spp. distribution and speciation (Chen et al. 2012; Lo and Wang 2021), and our findings further support and clarify these divergence processes.

Our results also explain why the jelly fig-*W.* sp. 3 currently inhabits altitudes between 500 and 2,000 meters in Taiwan; that is because their ancestors were adapted to an environment 6–7°C cooler during the Last Glacial Period (Lin 1963). Previous studies have shown that *W.* sp. 3 survived longer at lower temperatures and displayed positive selective signals in mitochondrial markers in contrast to *W.* sp. 2 (Wang et al 2013). In this study, we detected genome-wide signatures of positive selection related to energy production and metabolism in *W.* sp. 3, thereby providing genetic mechanisms for cold adaptations. Taken together, both demographic and selection analyses support the evolutionary scenario that the isolation of jelly fig-*W.* sp. 3 from mainland creeping fig-*W.* sp. 2 during the Last Glacial Period led to local adaptation to a colder environment, eventually manifesting as niche partitioning under the Holocene climate.

In addition to different habitat preferences, the two host varieties also exhibit divergent life history traits. Depending on geographical locations, male creeping figs produce three or four rounds of syconium per year (Ho et al. 1998), whereas, male jelly figs only produce one or two rounds (Shih 1988). Each round of male syconia is typically synchronized in a way such that they are ready to rear the offspring of fig wasps from the previous round. The life cycle of fig wasps is therefore completely dependent and synchronized with the host phenology. The generation times are thus different between the two *Wiebesia* species.

We observed a significantly higher substitution rate in *W.* sp. 2 than that of *W.* sp. 3 (Table 3), which is expected given the differences in generation time (Li et al. 1996). Apart from the effect of generation time, warmer temperature in lowland habitats may also increase mutation rate in ectothermic insects (Waldvogel, 2021). The substitution rate is even higher in *W.* sp. 2 from China (Table 3). The difference of substitution rates between *W.* sp. 2 populations across the Taiwan Strait is likely an effect of gene flow between *W.* sp. 2 and sp. 3 in Taiwan, as shown in demographic analysis.

### Genetic basis of host specificity in the two *Wiebesia* species

Olfactory coadaptation is arguably the single most important trait for the wasps in maintaining host-pollinator specificity in fig-fig wasp mutualism (Janzen 1979; Hossaert-McKey et al. 2010). Previous studies have shown that the two *F. pumila* varieties have different floral volatile profiles (Chen and Wu 2010; Chen et al. 2016). Therefore, it is reasonable to expect differentiation in chemosensory genes between *Wiebesia* species. Our gene expression and evolutionary analyses revealed that chemosensory genes are differentially expressed between *Wiebesia* species (Table 6) and exhibit signs of adaptive evolution (Table S7), suggesting their contribution in shaping host specificity. The behavioral experiments demonstrated host specificity in *W.* sp. 2 and *W.* sp. 3, although *W.* sp. 3 can also respond to the male syconia of creeping fig, a phenomenon that was noted also in field sampling (Chen et al. 2012; Wang et al. 2013). These findings highlight the role of olfactory adaptations in driving the evolutionary divergence of pollinators in response to host-specific chemical cues.

Among the chemosensory genes, *OBP11* is an interesting candidate that may explain the phenotypic divergence between the two wasps. We found a significantly higher expression of *OBP11* in *W.* sp. 2 compare to *W.* sp. 3. *Wiebesia* sp. 2 uses *OBP11* to detect the repellent molecule nonanal emitted from creeping fig (Wang et al. 2021). Receptive stage male syconia of jelly fig (the host of *W.* sp. 3) emitted relatively higher amount of nonanal than that of creeping fig (Chen and Wu 2010; Chen et al. 2016). We therefore speculate that the higher expression of *OBP11* in *W.* sp. 2 leads to a higher sensitivity to the repelling signal of nonanal than in *W.* sp. 3. This may explain why *W.* sp. 2 is less attracted by jelly fig; while *W*. sp. 3, though showing a preference, is more flexible in its attraction to both varieties.

### Dynamics of ORs and implications on long-term fig-fig wasp coevolution

The mass reduction in chemosensory receptor gene numbers among fig wasps has been attributed as a consequence of their mutualistic association with fig trees (Xiao et al. 2013; Xiao et al. 2021). On the other hand, a recent transcriptome study has demonstrated that while the total number of olfactory receptor (OR) genes is conserved and highly reduced in fig wasps, distinct lineages exhibit varied patterns of gains and losses (Yu et al. 2023). Our genome-based analysis confirmed this finding. Additionally, we showed that many novel OR genes arose from local tandem duplications, with these duplicated ORs exhibiting a higher frequency of positive selection signatures compared to copy-number-conserved counterparts, indicating functional divergence of duplicate genes. Expansion of chemosensory receptors has been linked to various olfaction-related adaptations in insects (Engsontia et al. 2014; Harrison et al. 2018). A famous example in hymenoptera is the independent expansions of the OR 9-exon subfamily in several eusocial lineages (Engsontia et al. 2015; Brand and Ramírez 2017; Legan et al. 2021). Thus, adaptive gene duplication in ORs, such as the one observed in subfamily Z, may have played a crucial role in the coevolution of fig floral scents and wasp olfactory recognition.

## Conclusion

Through multiomic and behavioral analysis of pollinating wasps, we detected genome-wide signatures of positive selection related to energy production and metabolism in *W.* sp. 3, suggesting potential adaptations to colder, mid-altitude environments. We also observed a significantly lower substitution rate in *W.* sp. 3 compared to *W.* sp. 2, consistent with the longer generation time associated with their high-altitude, slower-growing host, the jelly fig. Furthermore, the divergence in chemosensory gene expressions and adaptive duplications in olfactory receptors suggest mechanisms of coadaptation to host floral scents. Taken together, these findings illustrate that because the life cycles of fig hosts and their pollinating wasps are tightly linked, the life history traits and habitat preferences of fig hosts can directly impact the evolutionary processes of their associated pollinators, leaving distinct molecular signatures in their genomes. These results contribute to the broader understanding of coevolutionary dynamics, uncovering the genomic processes of codiversification.

Future studies should focus on exploring the full cophylogeny, including the two host varieties, as well as other copollinators such as *W.* sp. 1 (Chen et al. 2012). While we have performed multiomic and behavioral analysis, functional genomic studies and experimental validations incorporating data from wasps and figs are necessary to bridge the gap between molecular and gross phenotypes.

## Materials and Methods

### Biological materials and sequencing

Genomic DNA was extracted from 30 pooled individuals of each species using Purgene kit (Qiagen, USA); Whole body RNA from five stages (adult female, adult male, pupal female, pupal male, and larva) was extracted using TRIzol (ThermoFisher, USA). Library preparation and sequencing were performed by Novogene, Beijing, China. Long read sequencing of *W.* sp. 3 was done on a PacBio Sequel platform. DNA and RNA libraries that had insert size around 300-500bp were prepared for *W.* sp. 3 and *W.* sp. 2 and sequenced on Illumina Hiseq 4000 platform. The libraries and sample information are summarized in Table S1.

### Genome assembly and annotation

Short genomic reads were quality trimmed by Trimmomatic v0.36 (Bolger et al. 2014). Quality assessment was done by FastQC v0.10.1 (Andrews 2010). Jellyfish v2.2.6 (Marçais and Kingsford 2011) was used to estimate genome size from kmer. An initial genome was assembled from long reads by wtdbg v1.1.006 (Ruan and Li 2020) and evaluated by blobtools v1.1 (Laetsch and Blaxter 2017). Both short and long reads were filtered by blobtools to remove contaminations. Filtered long reads were assembled by wtdbg, and polished with arrow and pilon v1.2 (Walker et al. 2014). By the time of the analysis, the *W.* sp. 2 genome from population in China hasn’t been released. Therefore, the assembled *W* sp. 3 genome served as a backbone for the reference-based assembly of *W.* sp. 2. Short reads of *W.* sp. 2 were mapped against *W.* sp. 3 genome using BWA mem (Li and Durbin 2009). Variants from alignment file were called using samtools v1.2 (Li 2011). Consensus sequence was generated by bcftools v1.3 (Li 2011) in genomic regions that have a coverage between 1/3 to 2 times the mean coverage; regions that have coverage outside the range were substituted by Ns. Pilon was applied iteratively on the *W.* sp. 2 genome to update the genome based on information from short reads. Quast v4.6.3 (Gurevich et al. 2013) was used to calculate assembly statistics; busco v3 (Simão et al. 2015) was used to evaluate the completeness based on Hymenoptera odb9 dataset (Zdobnov et al. 2017). Foreign contigs detected by blobtools were excluded before annotation. D-Genies server (Cabanettes and Klopp 2018) was used to perform dotplot analysis on the assembled *W.* sp. 3 genome and the published *W.* sp. 2 (China) genome.

The two assembled genomes were annotated using the same pipeline. Annotation of repetitive DNAs based on homology was done by RepeatMasker v4.0 (Tarailo-Graovac and Chen 2009) with data from Repbase (Bao et al. 2015) and the ones predicted by RepeatModeler v1.0.1 (Tarailo-Graovac and Chen 2009). Repeat-masked genomes were created by bamtools v2.5.1 (Barnett et al. 2011), and were used in *ab initio* gene predictions. Gene predictions based on transcriptome, protein homology, and *ab initio* methods were loaded to Evidencemodeler v1.1.1 (EVM) (Haas et al. 2008) to generate consensus gene sets. Consensus gene sets were manually curated using Web Apollo v2.1.0 (Lee et al. 2013). Blast2GO v5.2.1 (Gotz et al. 2008), which integrated blast and interproscan (Jones et al. 2014) annotation results were used for functional and gene ontology (GO) annotation. Online server of BlastKOALA (Kanehisa et al. 2016) was used to map the gene sets to KEGG database (Kanehisa and Goto 2000).

### Demographic analysis

PPalign module in the PoolParty pipeline was used to call SNPs from short read libraries (Micheletti and Narum 2018). Reads were first trimmed by BBduk to remove adapters, contaminants, low-quality reads (< 20), and reads less than 25 bp in length (Bushnell 2014). Trimmed reads were aligned to *W.* sp3 assembly with a minimum mapping quality of 20 using the BWA-MEM (Li & Durbin, 2009). Duplicate reads were removed by SAMBLASTER (Faust and Hall 2014) and reads with mapping quality less than five were filtered in SAMtools before being converted to bam files (Li, 2011). SNPs were called in BCFtools with a SNP quality threshold of 20. After applying the initial filter criteria, the variant data set was limited to biallelic sites using VCFtools (Danecek et al. 2011).

The folded site frequency spectrum (SFS) for all populations were generated from the final variant calls (vcf files) using easy-SFS (https://github.com/isaacovercast/easySFS) (Gutenkunst et al. 2009). Demographic parameters were estimated from the joint SFS using coalescent simulations (fastsimcoal2) (Excoffier et al. 2021). The likelihood of the observed SFS under six complex demographic scenarios were calculated. Each model was run 100 replicated times considering 100 000–250 000 simulations for the calculation of the composite likelihood. We used an information-theoretic model selection approach based on the Akaike’s information criterion (AIC) to determine the probability of each model given the observed data. The mutation rate of 2.8 x 10^-8^ per site per year for fig wasp was applied (Xu et al. 2021).

### Phylogeny of studied fig wasp

The published transcriptome reads of *El. stueckenbergi* (SRR1502962) were *de novo* assembled with Trinity v2.8.4 (Haas et al. 2013). Transcript isoforms were filtered so that only the longest isoform was left as a representative of each gene. Protein sequences of *C. solmsi*, *Eu. verticillata* and *N. vitripennis* (Pteromalidae) were downloaded from publications (Zhou et al. 2015; Zhang et al. 2020). A total of 2684 busco genes that were single copy in all six species were used to reconstruct phylogeny of fig wasps. The genes were aligned with MAFFT v6 (Katoh and Standley 2013), filtered with trimal 2 (Capella-Gutiérrez et al. 2009) using default parameters, and then concatenated. Maximum likelihood tree of the concatenated busco genes was constructed using raxml-ng (Stamatakis 2014) with 100 bootstraps.

### Evolutionary rates and signatures of selection between *Wiebesia* species

Orthology between the four protein coding gene annotations: *W.* sp. 3, *W.* sp. 2 (Taiwan), *W.* sp. 2 (China), and *C. solmsi*, were determined using proteinortho v5.16 (Lechner et al. 2011). One to one orthologs were aligned by codon using PRANK v.14 (Löytynoja 2014), and filtered with Gblocks 0.91b (Castresana 2000). dN /dS ratio of each branch was estimated using codeml free ratio model from paml v4.9 (Yang 2007). GO enrichment analysis using Blast2GO were performed on the sets of genes with a dN /dS ratio larger than 1 with a FDR cutoff 0.05 (Benjamini and Hochberg 1995). To identify genes putatively under positive selection, Tajima’s D values were calculated using vcf-kit (Cook and Andersen 2017) on the biallelic SNPs of each species. A sliding window approach was adopted, with window and step sizes set at 5000 bp and 500 bp, respectively. Genes for which the majority of overlapping windows had negative Tajima’s D values were considered to be under positive selection.

### Fig wasp host attraction experiment

Eight combinations of lineage and sex were used for behavioral experiments (two wasp species × two syconia sexes × two host varieties). Each combination has a minimum of three experimental replicates (Table 5). To avoid influence from wild wasps, all the syconia used for entry experiment were sealed with mesh bag in pre-floral phase. Fifteen freshly emerged female adult wasps were introduced inside the mesh bag of each syconium once the syconium reaches receptive phase (based on the gathering of wild wasps outside the mesh). The number of entry wasps were calculated 24 hours after the introduction. Entering rate is calculated as the number of entry wasps / 15. The *t* test was used for statistical testing between combinations.

### Cross species differential expression analysis of the *Wiebesia* species

Four biological replicates of adult female transcriptome were sequenced for each species. Transcript per million reads (TPM) for each gene was calculated with the Perl script “align_and_estimate_abundance.pl” provided by Trinity (Haas et al. 2013) for each library. Kallisto v0.44 (Bray et al. 2016) was used for abundance estimation. TPM values were adjusted by ortholog length using the ‘scaledTPM’ method from Bioconductor package ‘tximport’ (Soneson et al. 2016). Cross sample normalization was then applied on the adjusted TPM values using TMN method (Robinson et al. 2010). Biological replicates validation was done by principal component analysis and hierarchical clustering using R. Differential expression analysis was performed on all single copy orthologs that are expressed in at least two libraries using the bioconductor package edgeR (Robinson et al. 2010) with minimum log fold change set to 4 and a FDR cutoff of 0.05. GO enrichment analysis of differentially expressed genes was done by Blast2GO based on annotation of *W.* sp. 3, with a FDR cutoff of 0.05.

### Manual annotation of olfactory receptors and evolutionary analysis

Amino acid sequences of Hymenoptera ORs (Zhou et al. 2015) were downloaded as query dataset. Manual annotation of chemosensory genes in the four fig wasp species: *W.* sp. 3, *W.* sp. 2, *El. stueckenbergi*, *Eu. verticillata* were done with two approaches: by blasting the annotated gene sets against the query set, and by homology-based gene model prediction with GenBlastA v1.0.4 (She et al. 2009) and GeneWise v2.2 (Birney et al. 2004) based on query sequences. Results of the two approaches were manually combined under Web Apollo. Maximum likelihood phylogeny of fig wasp ORs was created by raxml-ng v1.0.2 and labeled using MEGA7 (Kumar et al. 2016). Orthology of genes was determined by both proteinortho and phylogenetic relationships. TMHMM v2.0 server (Krogh et al. 2001) was used to annotate transmembrane domain in the OR genes. Gene birth and death events were interpreted using Notung 2.9.1.5 (Chen et al. 2000) using raxml-ng tree as input. The rearrange threshold for bootstrap values was set to 70. Tandem repeats were identified by manual inspecting the coordinates of the OR genes.

Both *El. stueckenbergi* and *Eu. verticillata* were excluded in cds-based analysis because of annotation quality. OR gene tree was divided into two types of orthogroups. Each orthogroup was consisted of at least three sequence from at least two species. A simple orthogroup is consisted of single copy orthologs from each of the three fig wasp species only, while a complex orthogroup contains paralogs. Each orthogroup was aligned by codon using PRANK. Low quality sites were filtered by Gblocks. The filtered alignments were manually examined so that incomplete genes can be excluded. Branch-site test using aBSREL (Smith et al. 2015) from online server of Datamonkey (Weaver et al. 2018) was performed on all the subtrees to identify branches under positive selection. A custom R script was used to adjust for multiple testing using FDR, and perform Fisher exact test to test whether positive selection occurred significantly differently between groups. The significance cutoff for FDR and Fisher exact test p value are both 0.05.

## Supplementary Materials

Supplementary data can be found online.

## Author contributions

H. W. conceived the study. B. L. and H. W. designed the genomic study. B. L. collected the samples, assembled the genomes, performed the comparative genomic, transcriptomic, and OR evolutionary analyses. H. L. performed population genomic analyses. S. W. analyzed evolutionary rates. S. K. and W. W. designed and performed the behavioral experiments. X. L. and H. W. secured funding and supervised the project. B. L. and H. W. wrote the manuscript.

## Acknowledgments

This work was supported by Ministry of Science and Technology, Taiwan (105-2628-B-002 - 015 -MY3). We thank Yanling Wen for the help with bioinformatic analyses; Liduo Yin and Yen-Hua Yeh for sample processing; ShenShan Aiyu, SanShuiLi Aiyu, SinPing Farm, and Miaoli District Agricultural Research and Extension Station for providing fig wasp samples; Fiona Puntieri, Luo-Yu Chang, Mai Phan, Paolo Franchini, Shu-Miaw Chaw, Wen-Hao Tan, and Wen-Shi Guo for commenting on the manuscript.

## Data availability

Genomes and the raw reads used for this study can be found under the NCBI BioProject: PRJNA730931. Additional information including functional annotations, OR genes, and scripts for this study can be found https://github.com/JohnsonStev/Wiebesia_wasp_genomes

## Notes

https://github.com/JohnsonStev/Wiebesia_wasp_genomes

### Competing Interest Statement

The authors have declared no competing interest.

