## Supplementary Materials for "Genomes of *Wiebesia* fig wasps reveal the adaptation and codiversification in fig-fig wasp mutualism"

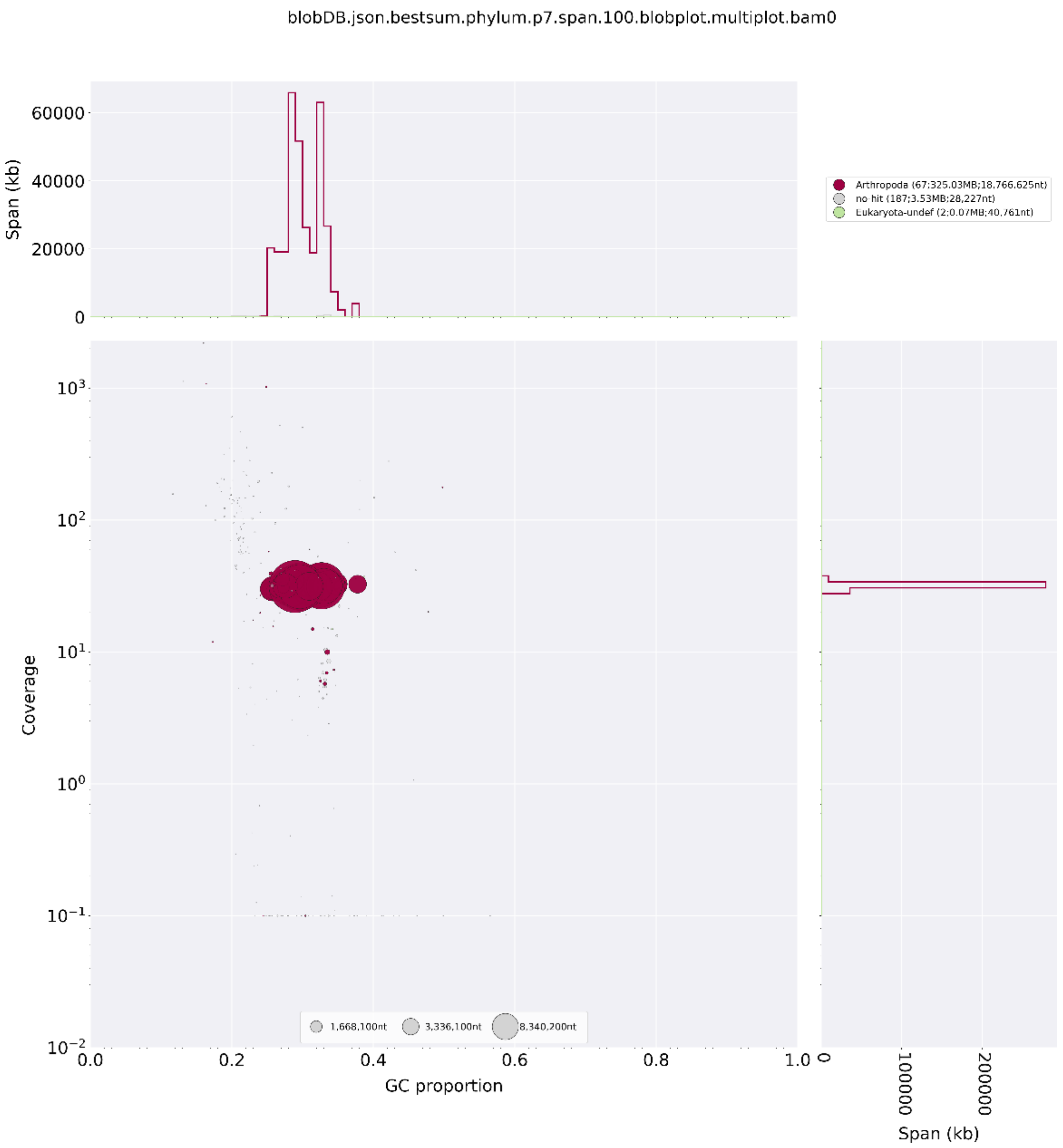

Fig. S1. Blobplot of *Wiebesia* sp. 3 genome after filtering. Colors represent the phylum each contig was assigned to. Most of the larger contigs have similar coverage and are assigned to Arthropoda (colored red).

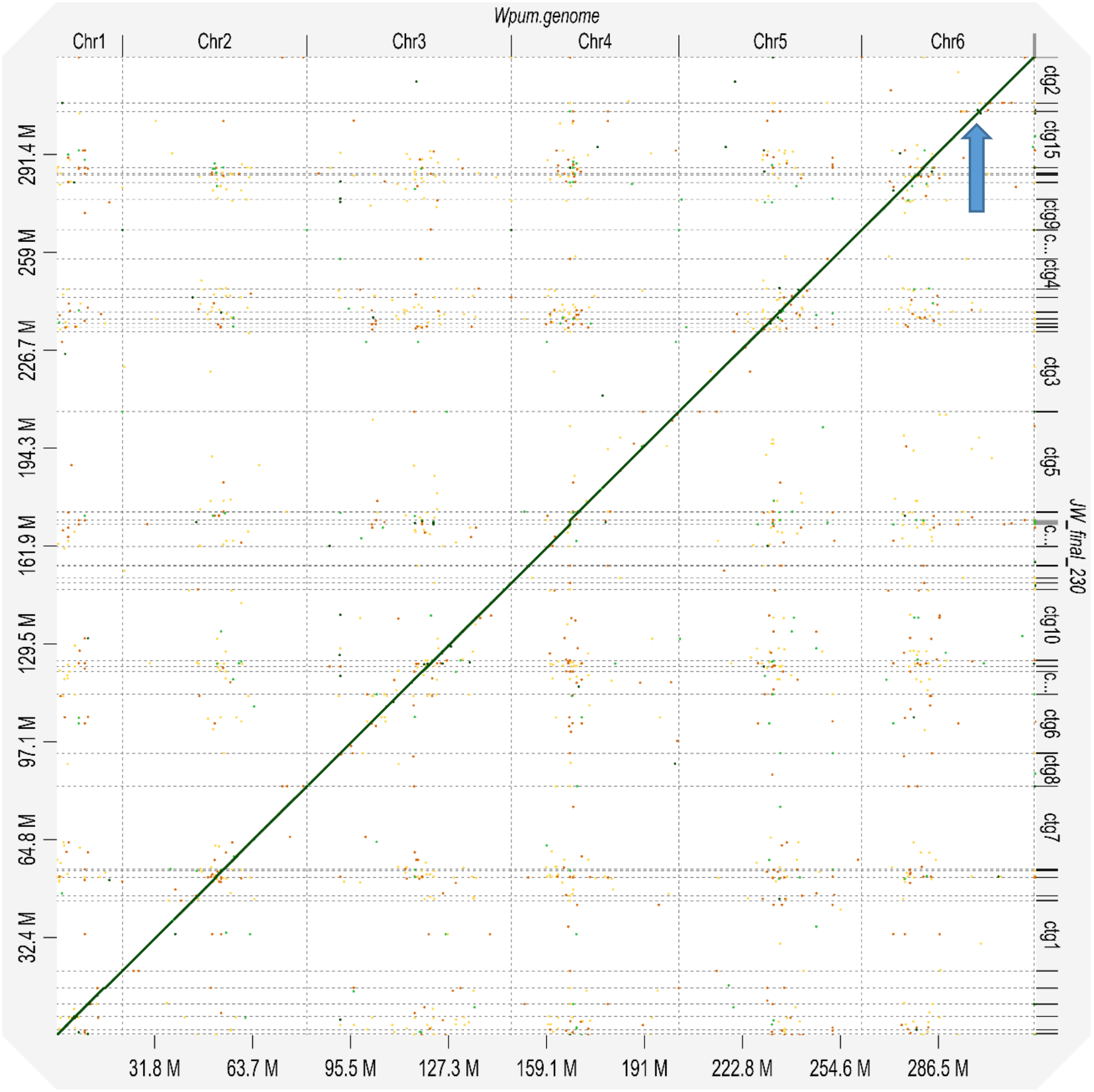

Fig. S2. Whole genome alignment dotplot of the *Wiebesia* spp. The published *W.* sp. 2 genome from China (x axis) and the newly sequenced *W.* sp. 3 (y axis) are conserved in synteny. Only an inversion 0.9 Mb in size was identified on chromosome 6 of *W.* sp. 2. (blue arrow)

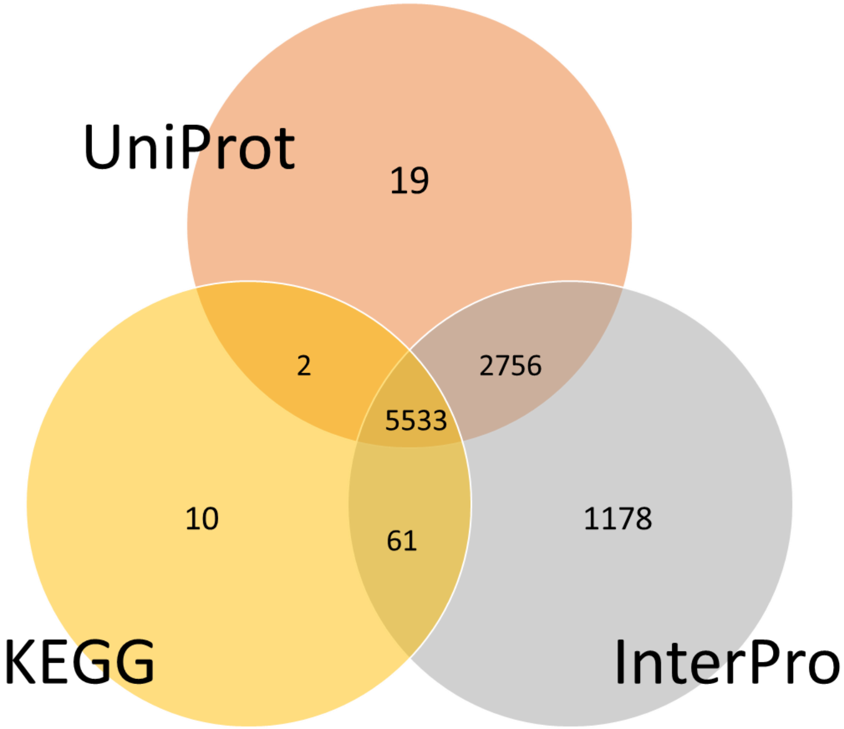

Fig. S3. Venn diagram showing functional annotation of *Wiebesia* sp. 3 genome. Numbers represent genes annotated based on each database.

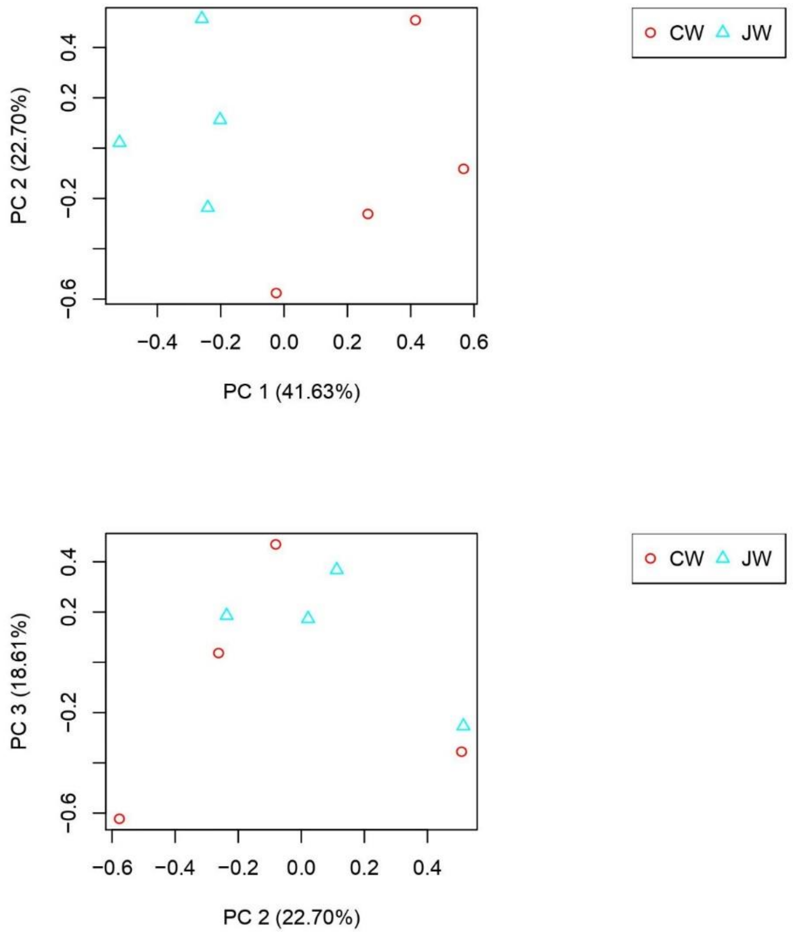

Fig. S4. Principal component analysis of normalized adult female expressional profiles. Note that the two species can be separated by the first principal component. CW: *W.* sp. 2; JW: *W.* sp. 3

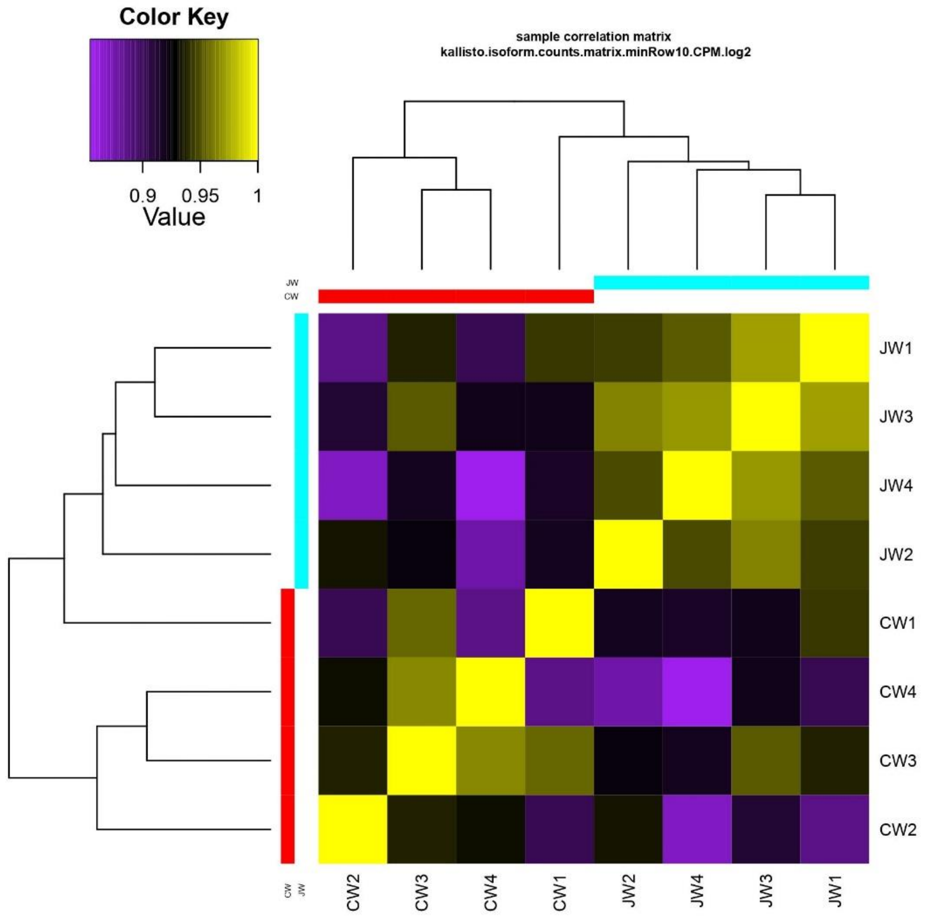

Fig. S5. Correlation matrix of normalized adult female expression profiles. CW: *W.* sp. 2; JW: *W.* sp. 3

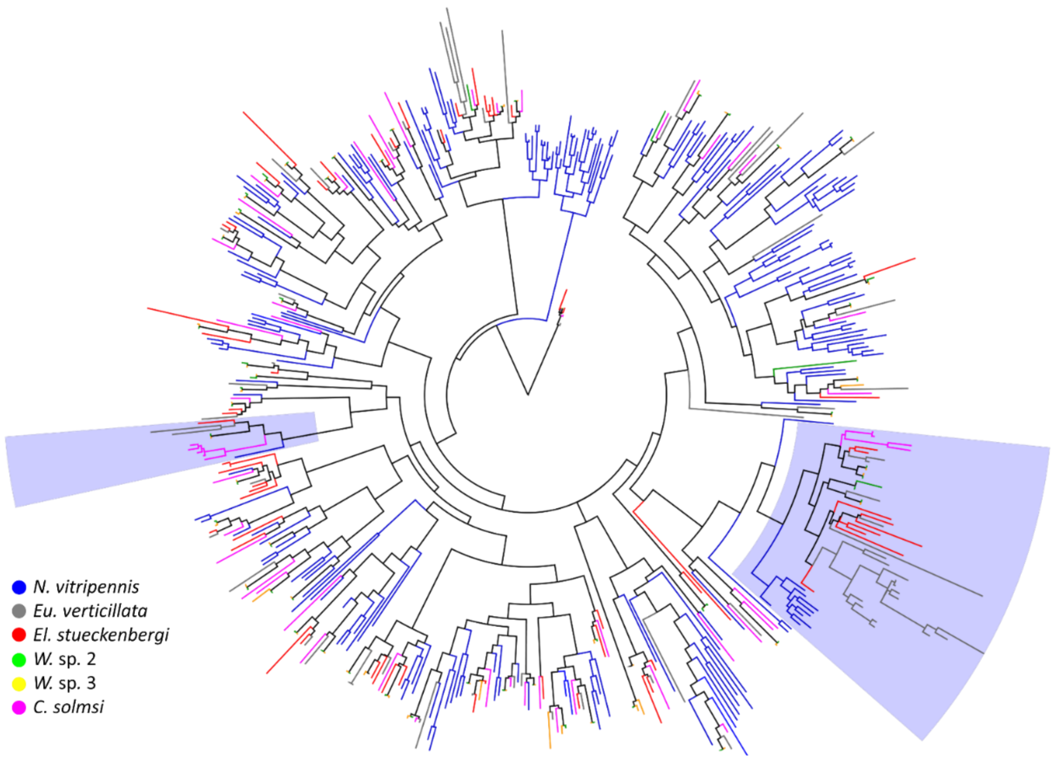

Fig. S6. Maximum likelihood phylogeny of olfactory receptors. Five fig wasps and an outgroup parasitoid wasp species *Nasonia vitripennis* were included. The tree is rooted by the Orco genes. The two shaded regions were detailed in Fig 4, demonstrating examples of lineage-specific expansions.

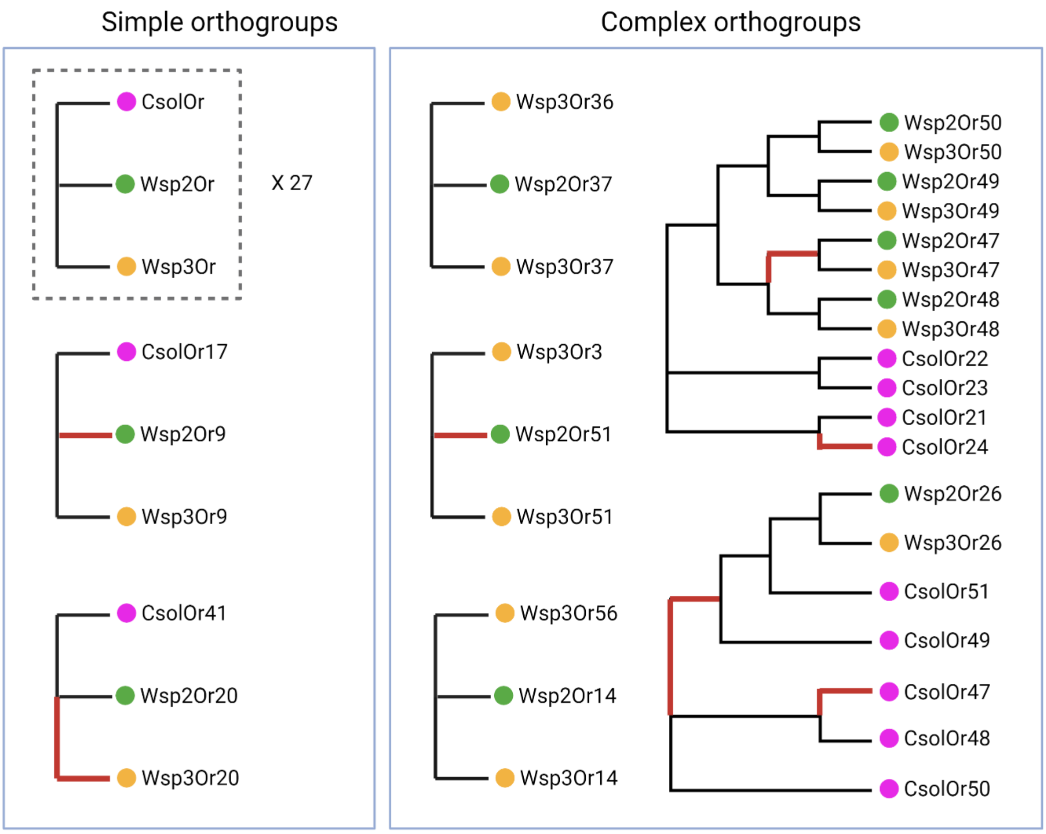

Fig. S7. Simple orthogroups and complex orthogroups of olfactory receptors. Branches that show signatures of selection were marked in red. Created with bioRender.com.

Table S1. Information of sequenced libraries.

| Library name | Description | Collection site | Sequenced bps |
|---|---|---|---|
| **JW_DNA_long_reads** | Adult male | Pingtung, Taiwan | 28.0G |
| **JW_DNA** | Adult male | Pingtung Taiwan | 22.8G |
| **CW_DNA** | Adult female | Chiayi, Taiwan | 27.8G |
| JW_RNA_AF1 | Adult female | Nantou, Taiwan | 7.6G |
| JW_RNA_AF2 | Adult female | Nantou, Taiwan | 14.5G |
| JW_RNA_AF3 | Adult female | Nantou, Taiwan | 6.2G |
| JW_RNA_AF4 | Adult female | Nantou, Taiwan | 6.8G |
| JW_RNA_AM | Adult male | Nantou, Taiwan | 7.8G |
| JW_RNA_PF | Pupa female | Nantou, Taiwan | 9.9G |
| JW_RNA_PM | Pupa male | Nantou, Taiwan | 8.5G |
| JW_RNA_L | Larva both sex | Nantou, Taiwan | 3.2G |
| CW_RNA_AF1 | Adult female | Matsu, Taiwan | 7.4G |
| CW_RNA_AF2 | Adult female | Matsu, Taiwan | 15.7G |
| CW_RNA_AF3 | Adult female | Chiayi, Taiwan | 6.7G |
| CW_RNA_AF4 | Adult female | Chiayi, Taiwan | 5.3G |
| CW_RNA_AM | Adult male | Matsu, Taiwan | 6.3G |
| CW_RNA_PF | Pupa female | Matsu, Taiwan | 7.6G |
| CW_RNA_PM | Pupa male | Matsu, Taiwan | 7.7G |
| CW_RNA_L | Larva both sex | Hualien, Taiwan | 4.2G |

JW=jelly fig wasp; CW= creeping fig wasp.

Table S2. Statistics and annotation quality of fig wasp sequencing projects

| Species | Sequencing platform | Assembly level | Scaffolds | Total size (bp) | N50 (bp) | Gene count | BUSCO (Hymenoptera gene set) |
|---|---|---|---|---|---|---|---|
| *Ceratosolen solmsi** | Illumina | Scaffold | 7397 | 278M | 9.5M | 11412 | 93.9% |
| *Eupristina verticillata*** | PacBio | Scaffold | 768 | 387 M | 3.1M | 14312 | 89.7% |
| *Wiebesia. pumilae**** (sp. 2) | PacBio | chromosome | 102 | 318M | 10.9M | 12316 | 91.3% |
| Assembled in this study | | | | | | | |
| *Wiebesia* sp. 3 | PacBio | Scaffold | 230 | 324M | 18.6M | 10072 | 95.5% |
| *Wiebesia* sp. 2 | Illumina | Scaffold (mapping-based) | - | - | - | 10037 | 95.7% |
| *Elisabethiella stueckenbergi* | Illumina | *de novo* transcriptome | - | 59.7M | 1746 | 39697 | 79.70% |

*(Xiao et al. 2013)
**(Zhang et al. 2020)
***(Wang et al. 2021)

Table S2 Repetitive DNA contents in *Wiebesia* sp. 3.

| | **Repeatmodeler** | **Repbase** | **RepeatProtein-Mask** | **Combined** | **Percentage** |
|---|---|---|---|---|---|
| DNA | 3852826 | 1459065 | 224966 | 4711271 | 1.45% |
| LINE | 722260 | 145464 | 165194 | 897960 | 0.28% |
| LTR | 1524146 | 949446 | 3171361 | 4175138 | 1.29% |
| Low complexity | 2048667 | 2121798 | 2190975 | 2220568 | 0.69% |
| Other | | | 246 | 246 | 0.00% |
| SINE | 71052 | | | 71052 | 0.02% |
| Simple repeat | 9213329 | 9485476 | 9743499 | 9848319 | 3.04% |
| Unknown | 11554309 | 28810 | | 11583026 | 3.58% |
| rRNA | | 29902 | | 29902 | 0.01% |
| tRNA | | 2204 | | 2204 | 0.00% |
| Total | 28986589 | 14222165 | 15496241 | 33539686 | 10.36% |

DNA: DNA transposons; LINE: long interspersed nuclear elements; LTR: long terminal repeats; SINE: short interspersed nuclear elements.

Table S3. *Wiebesia* sp. 3 gene models predicted by each software.

| **Gene prediction software/pipeline** | **Predicted genes** | **BUSCO results*** |
|---|---|---|
| hisat2+stringtie+transdecoder | 20162 | 90.1% |
| GeneWise/*N. vitripennis* | 20313 | 82.1% |
| GeneWise/*A. mellifera* | 15544 | 84.4% |
| GeneWise/*C. solmsi* | 18510 | 91.0% |
| GeneWise/*C. floridum* | 22633 | 88.1% |
| GeneWise/*T. pretiosum* | 23246 | 83.4% |
| SNAP (trained by *C. solmsi*) | 73718 | 43.6% |
| GlimmerHMM (trained by *C. solmsi*) | 28149 | 26.2% |
| Augustus (trained by BUSCO) | 31374 | 71.7% |
| GeneID | 15300 | 54.4% |
| FgenesH | 48716 | 48.8% |
| EVM | 13342 | 93.0% |
| Apollo (manually curated) | 10072 | 95.5% |

*BUSCO based on hymenoptera odb9 dataset.

Table S4 Parameters inferred from coalescent simulations with Fastsimcoal2 under different models

| **Model** | **No gene flow** | **Early gene flow** | **Constant gene flow** | **Recent gene flow** | **Early-recent gene flow** |
|---|---|---|---|---|---|
| **AIC** | 16,274,278 | 16,230,181 | 16,282,779 | 15,663,005 | 15,652,928 |
| **$Ne_{\_sp2}$** | 309,261 | 345,113 | 319,653 | 324,733 | 329,295 |
| **$Ne_{\_sp3}$** | 260,620 | 264,189 | 267,284 | 257,984 | 152,572 |
| **$T_{\_Div}$** | 497,188 | 539,714 | 517,851 | 529,503 | 547,178 |
| **$ME_{2\rightarrow3}$** | | 7.81E-09 | 1.10E-09 | | 1.85E-09 |
| **$ME_{3\rightarrow2}$** | | 1.54E-08 | 2.53E-09 | | 1.61E-07 |
| **$T_{\_Stop}$** | | 250,796 | | | 80,752 |
| **$T_{\_Mig}$** | | | | 9,609 | 11,243 |
| **$MR_{2\rightarrow3}$** | | | | 1.14E-08 | 1.04E-08 |
| **$MR_{3\rightarrow2}$** | | | | 1.43E-09 | 2.04E-09 |

Ne: Effective population size

$T_{\_Div}$: Time of divergence in years

ME: Early migration rate

$T_{\_Stop}$: Time of early migration stop in years

$T_{\_Mig}$: Time of recent migration resumed in years

MR: Recent migration rate

Table S5 Gene ontologies enriched for genes with negative Tajima's D in *Wiebesia* sp.3. All GOs significant at 0.05 FDR level were shown

| GO term | GO ID | FDR |
|---|---|---|
| **Process** | | |
| Nucleobase-containing compound biosynthetic process | 0034654 | 0.030 |
| Monoatomic ion transport | 0006811 | 0.043 |
| Primary metabolic process | 0044238 | 0.043 |
| Cellular metabolic process | 0044237 | 0.012 |
| Nitrogen compound metabolic process | 0006807 | 0.043 |
| **Component** | | |
| proteasome complex | 0000502 | 0.043 |
| **Function** | | |
| Organic cyclic compound binding | 0097159 | 0.005 |
| Protein binding | 0005515 | 0.043 |
| Heterocyclic compound binding | 1901363 | 0.005 |

Table S6 Chemosensory genes that are differentially expressed between females of *Wiebesia* sp. 2 and sp. 3.

| **Gene ID (*W.* sp. 3)** | **Up regulated** | **Log fold change** |
|---|---|---|
| **OR** | | |
| WAW_00001067 | *W.* sp 2 | 4.42 |
| WAW_00002230 | *W.* sp 2 | 2.07 |
| WAW_00002856 | *W.* sp 3 | 2.48 |
| WAW_00003378 | *W.* sp 3 | 3.19 |
| WAW_00006086 | *W.* sp 2 | 5.18 |
| WAW_00006388 | *W.* sp 2 | 2.15 |
| WAW_00006640 | *W.* sp 2 | 5.43 |
| **GR** | | |
| WAW_00004441 | *W.* sp 3 | 3.52 |
| WAW_00006236 | *W.* sp 2 | 3.03 |
| **IR** | | |
| WAW_00002511 | *W.* sp 2 | 3.54 |
| WAW_00008616 | *W.* sp 2 | 2.23 |
| WAW_00009024 | *W.* sp 2 | 2.82 |
| **CSP** | | |
| WAW_00005774 | *W.* sp 2 | 5.09 |
| WAW_00005775 | *W.* sp 2 | 3.10 |
| **OBP** | | |
| WAW_00001780* | *W.* sp 2 | 2.67 |
| WAW_00008197 | *W.* sp 3 | 4.90 |
| WAW_00009356 | *W.* sp 2 | 2.56 |

*WAW_00001780=*OBP11*; Genome annotations can be found in the github page.

Table S7 Patterns of selection in chemosensory genes of *Wiebesia* spp.

| | **ORs** | **GRs** | **IRs** | **OBPs** | **CSPs** | **NCs*** |
|---|---|---|---|---|---|---|
| Single-copy in *Wiebesia* | 56 | 4 | 10 | 12 | 8 | 9141 |
| Differentially expressed | 7** | 2** | 3** | 2 | 3** | 415 |
| dN/dS>1 | 9** | 1 | 0 | 1 | 0 | 632 |
| average dN/dS | 0.4248 | 0.3826 | 0.2704 | 0.1088 | 0.0966 | 0.2315 |
| average dN | 0.0063 | 0.0051 | 0.0039 | 0.0018 | 0.0007 | 0.0028 |
| average dS | 0.0148 | 0.0134 | 0.0146 | 0.0168 | 0.007 | 0.0119 |
| Single-copy in *Wiebesia* and *C. solmsi* | 34 | 4 | 10 | 9 | 8 | 8182 |

*Non-chemosensory genes

** Significantly enriched versus background NC genes, Fisher exact test $p < 0.05$

Table S8 Average number of transmembrane domain (TD) in ORs in the studied fig wasps and fruit fly. Species with average TD numbers above 5 were included in branch test anaylsis.

| **Species** | **Mean number of TDs (standard deviation)** |
|---|---|
| *W.* sp. 2* | 5.50 (2.05) |
| *W.* sp. 3* | 5.32 (1.76) |
| *El. verticillata* | 3.85 (2.55) |
| *Eu. stueckenbergi* | 1.57 (1.38) |
| *C. solmsi** | 5.85 (1.18) |
| *D. melanogaster* | 5.72 (1.12) |

*Species included in branch site selection test.

### Supplementary Material and Methods

**Biological Materials and Species Validation**

Male jelly fig syconia were collected from Pingtung, Taiwan (22.45° N, 120.43° E, 650 m), and male creeping fig syconia were obtained from Chiayi, Taiwan (23.33° N, 120.28° E, 55 m). Wasps collected from a single syconium were pooled, washed with distilled water, and stored in ethanol (EtOH) prior to DNA extraction. Transcriptomes from whole bodies at five different life stages for each species were sequenced for gene prediction: (1) adult female, (2) adult male, (3) pupal female, (4) pupal male, and (5) larva. Additionally, three adult female samples per species were sequenced as biological replicates for cross-species differential expression analysis. Previous studies have reported that creeping fig syconia may occasionally contain *Wiebesia* sp. 3 individuals (Wang et al. 2013). However, the collection site for creeping fig syconia in this study was distant from natural jelly fig habitats, reducing the likelihood of pooling individuals from different species. To confirm species purity, short reads from creeping fig wasps were mapped to a published cytochrome c oxidase subunit I barcode sequence (accession: JN184048.1) using BWA v0.7.10-r789 mem. The alignment was assessed manually using the Integrative Genomics Viewer (IGV) (Robinson et al. 2011) to ensure the reads matched the characteristics of *W.* sp. 2.

**Genome Assembly and Contamination Filtering**

Raw Illumina paired-end (PE) reads from genomic sequencing of both species were quality-trimmed using Trimmomatic v0.36. Reads were trimmed at both ends if their quality score was below 3. Additionally, reads with an average quality below 15 in a 4-base sliding window were cut, and reads shorter than 36 bp were discarded. Long reads of *W.* sp. 3 were assembled into contigs using wtdbg v1.1.006 with parameters -k 0 -p 21 -S 4 --edge-min 4. This initial assembly was polished using arrow and pilon v1.2. The level and origin of non-fig wasp DNA in this assembly were evaluated with Blobtools v1.1, which annotated contigs by blasting them against the NCBI nucleotide database using the parameter --x bestsum. Based on the blobplot, all reads mapping to contigs identified as Arthropods were retained. Additionally, reads mapping to contigs with GC content < 0.4 and mean coverage > 13 were retained for assembly. The filtered reads were assembled using the same parameters as the initial assembly.

**Genome Annotation Pipeline**

Repetitive DNA annotation based on homology was conducted using RepeatMasker v4.0, with input comprising Hymenoptera sets from Repbase and predictions by RepeatModeler v1.0.1. Transposable elements were identified using RepeatProteinMask from RepeatMasker. Protein-coding genes were predicted using multiple forms of evidence. For transcriptome-based prediction, RNA-seq reads from five life stages were aligned to the genome using HISAT2 v2.1 (Kim et al. 2015). Each library was assembled independently using StringTie v1.3 (Pertea et al. 2015), and the assembled transcripts were merged using StringTie "--merge". Transdecoder v5.3 (Haas et al. 2013) was used to predict ORFs of transcripts, with "retain_blastp_hits" taking blastp results from the UniProt database (UniProt Consortium 2012). For homologous gene prediction, gene sets from five Hymenoptera species—*Ceratosolen solmsi* (Xiao et al. 2013), *Nasonia vitripennis* (Werren et al. 2010), *Copidosoma floridanum* (i5K Consortium 2013), *Trichogramma pretiosum* (Lindsey et al. 2018), and *Apis mellifera* (Consortium 2006)—were procured from NCBI ftp. A non-redundant gene set for each species was created by selecting the longest isoform as the representative of each gene using a custom Perl script. The genes were aligned to the assembly with GenBlastA v1.0.4, and aligned regions, along with 3 Kbp flanks, were used in GeneWise v2.2 (Birney et al. 2004) predictions. For *ab initio* predictions, AUGUSTUS v3.3.1 (Stanke and Morgenstern 2005), SNAP (Korf 2004), geneid v1.3 (Blanco et al. 2007), the FGENESH server (Solovyev et al. 2006), and GlimmerHMM v3.0.1 (Majoros et al. 2004) were applied to repeat-masked assemblies. AUGUSTUS was run with parameters generated by the genome mode of BUSCO using the Hymenoptera odb9 dataset, incorporating RNA-seq-derived intron hints. Geneid and FGENESH used *Nasonia* parameters, whereas GlimmerHMM and SNAP were trained with the *C. solmsi* gene set. Protein alignments, RNA-seq alignments, Transdecoder-predicted ORFs, and *ab initio* predictions were input into EvidenceModeler v1.1.1 for consensus gene calling. Consensus gene sets were curated manually in Web Apollo v2.1.0, focusing on eliminating *ab initio*-only gene models and refining fragmented models.

**Manual Annotation of Chemosensory Genes**

Published sequences of olfactory receptors (OR) (Zhou et al. 2015), gustatory receptors (GR) (Zhou et al. 2015), chemosensory proteins (CSP) (Kulmuni and Havukainen 2013), odorant binding proteins (OBP) (Forêt and Maleszka 2006; Vieira et al. 2012), and ionotropic glutamate receptors (iGluRs) (Croset et al. 2010) were gathered as query datasets. Chemosensory genes were annotated through two approaches: homology-based gene prediction as previously described and BLASTp searches on both EVM-predicted and curated gene sets. Results from both methods were merged manually using Web Apollo. For ionotropic receptor (IR) annotation, iGluRs were aligned with query sequences using MAFFT 7.407 with parameters "--localpair --maxiterate 1000", followed by phylogenetic tree construction using RAxML 8.2.12 with parameters "-# 20 -m PROTGAMMAAUTO" and 1000 bootstrap iterations. Similarities to known IRs were identified by examining the phylogenetic tree.

**Evolutionary Rate Analysis**

Orthologs between *W.* sp. 2 and *W.* sp. 3 were identified by best BLAST reciprocal hits (BBRH) of amino acid sequences using ProteinOrtho v5.16. A total of 9,352 1:1 orthologs were aligned by codon using PRANK v.14, resulting in 9,218 sequences maintaining complete triplet ORFs. Alignments were filtered with Gblocks 0.91b, using parameters: -t=c -e=-gb1 -b4=6 -b5=a. dN/dS ratios for filtered alignments were calculated using pairwise Codeml from PAML 4.9 with the parameter: CodonFreq = 0. For genes with no synonymous substitutions (dS = 0), a genome-wide median dS was assigned. Alignments with dS > 5 were excluded to remove poor alignments or saturation artifacts. Lineage-specific dN/dS were computed for single-copy orthologs between *W.* sp. 2, *W.* sp. 3, and outgroup *C. solmsi* using a free-ratio model. To compare substitution rates between Taiwanese and Chinese *W*. sp. 2 populations, a three-species free-ratio model was applied twice, each incorporating a different W. sp. 2 population.

**Fig Wasp Host Attraction Experiment**

For the host attraction experiments, female *W*. sp. 2 and creeping fig syconia were sourced from Taipei, Taiwan (25.01° N, 121.54° E, 14 m), while female *W*. sp. 3 and jelly fig syconia were collected from Nantou, Taiwan (24.10° N, 120.86° E, 735 m). A total of eight experimental settings were performed, varying by wasp species, syconia sex, and host variety, with multiple replicates per condition (details in Table 5). Fifteen freshly emerged adult female wasps were introduced to a mesh bag enclosing each syconium at its receptive phase, identified by observing wild wasps aggregating outside the mesh bag. All experiments were conducted between 9:00 and 13:00, encompassing the typical period of heightened wasp activity (personal observation).
